# The consequences of lipid remodelling of adipocyte cell membranes being functionally distinct from global lipid storage during obesity

**DOI:** 10.1101/712976

**Authors:** Ke-di Liu, Animesh Acharjee, Christine Hinz, Sonia Liggi, Antonio Murgia, Julia Denes, Melanie K Gulston, Xinzhu Wang, Yajing Chu, James A. West, Robert C Glen, Lee D. Roberts, Andrew J. Murray, Julian L. Griffin

## Abstract

Obesity is a complex disorder where the genome interacts with diet and environmental factors to ultimately influence body mass, composition and shape. Numerous studies have investigated how bulk lipid metabolism of adipose tissue changes with obesity, and in particular how the composition of triglycerides (TGs) changes with increased adipocyte expansion. However, reflecting the analytical challenge posed by examining non-TG lipids in extracts dominated by TGs, the glycerophospholipid (PL) composition of cell membranes has been seldom investigated. PLs contribute to a variety of cellular processes including maintaining organelle functionality, providing an optimised environment for membrane-associated proteins and as pools for metabolites (e.g. choline for one-carbon metabolism and for methylation of DNA). We have conducted a comprehensive lipidomic study of white adipose tissue in mice who become obese either through genetic modification (*ob/ob*), diet (high fat diet) or a combination of the two using both solid phase extraction and ion mobility to increase coverage of the lipidome. Composition changes in seven classes of lipid (free fatty acids, diglycerides, TGs, phosphatidylcholines, lyso-phosphatidylcholines, phosphatidylethanolamines, and phosphatidylserines) correlated with perturbations in one-carbon metabolism and transcriptional changes in adipose tissue. We demonstrate that changes in TGs that dominate the overall lipid composition of white adipose tissue are distinct from diet-induced alterations of PLs, the predominant components of the cell membranes. PLs correlate better with transcriptional and one-carbon metabolism changes within the cell, suggesting the compositional changes that occur in cell membranes during adipocyte expansion have far-reaching functional consequences.

## Introduction

Adipose tissue is the largest energy storage organ in the body. The tissue also has a complex role in regulating whole-body metabolism, including being an endocrine organ secreting adipokines (e.g. leptin, adiponectin, IL-6, TNFα) (1) which impact on multiple functions including appetite and energy balance, and which contribute to systemic inflammation, insulin sensitivity and angiogenesis (2-4). Increased adiposity may result in the accumulation of toxic lipid species which impair normal cellular functions (termed lipotoxicity), innate immune responses in adipose tissue and systemic inflammation, which combined with ectopic fat storage produces dysfunction in the liver, muscle, gut and pancreas (5-7). Thus, how fat is stored in adipose tissue has an important impact on whole body metabolism.

However, it is not only the total fat storage capacity which has an impact on adipose tissue health, but also the lipid composition in terms of classes of lipids stored, and degree of unsaturation and chain length of these lipid species (8). Membrane phospholipids (PLs) typically contain a high proportion of saturated fatty acids which make the bilayer relatively rigid, in turn having consequences for protein function and membrane transport (9). Increased abundance of monounsaturated (MUFA) and polyunsaturated (PUFA) fatty acids within PLs can increase the fluidity of membranes, improving insulin sensitivity and the recruitment of proteins such as Glucose Transporter 4 (GluT4). Conversely, depletion of total ω-3 long chain PUFAs are associated with hepatic insulin resistance, enhanced sterol regulatory element-binding protein 1c (SREBP-1c) and decreased peroxisome proliferator-activated receptor alpha (PPARα) activity, favouring lipogenesis over fatty acid (FA) oxidation (10). The lipid composition of the cell membrane also has an important role in inflammation (11). For example, the interaction of ω-6 fatty acids with ω-3 fatty acids found in the cell membrane determines the production of lipid mediators, including eicosanoids, which have important consequences for inflammation (12).

There are three major origins for the FAs used as lipid building blocks: (i) lipids derived from the diet, (ii) *de novo* lipogenesis (DNL) in liver and adipose tissue whereby carbohydrates are converted to saturated FAs, and (iii) enzymatic modifications (desaturation, elongation and oxidation of the FA). SREBP1 and leptin have activating and inhibitory effects on the expression of lipogenic genes involved in the conversion of carbohydrates to FAs such as fatty acid synthase (FASN) and acetyl-CoA carboxylase (ACC) (13). In turn, elongation and desaturation of FAs are controlled by essential enzymes like long-chain fatty acid elongase 6 (ELOVL6) and stearoyl CoA desaturase-1 (SCD-1, Δ9 desaturase), both of which are also SREBP-1c targets (14). A change in the set point control of Insig1/SREBP1 facilitates lipogenesis and availability of appropriate levels of FA unsaturation through SCD-1 expression (9, 15).

Besides being influenced by hormones and adipokines, expression of lipid metabolising genes can also be altered through epigenetic mechanisms. Epigenetic and chromatin-modifying proteins have been found to contribute to adipogenesis and maintenance of mature adipocytes through PPARγ (16). Furthermore, methylation of the genes encoding PPARα and the glucocorticoid receptor have been shown to be decreased when maternal protein is restricted, with this mechanism having a long term impact on systemic metabolism (17). In addition, methylation of the gene encoding the melanocortin-r receptor is associated with long term exposure to a high fat diet (18). There are also important mechanistic and metabolic reasons to associate changes in the composition of diets to higher proportions of FA intake with epigenetic modifications of DNA. S-adenosyl methionine (SAM) is mainly derived from methionine via the enzyme methionine adenosyltransferase, an enzyme which is part of the choline and one-carbon (1-C) pathway and hence altered by increased consumption of phosphatidylcholines (PCs) (19-21). A sufficient SAM concentration is also required to establish the normal ratio of phosphatidylethanolamine (PE) to PC in the cell (22) providing another link between total PL metabolism and epigenetic modifications.

In this study we have investigated how metabolic composition (lipidome and metabolome) of adipose tissue is influenced in both diet and genetic induced models of obesity. We describe how adipose tissue attempts to maintain the composition of the PL fraction in the face of obesity, but ultimately this produces a range of physiological changes including altered membrane fluidity, altered transcriptional profiles and alterations in the methylation status of the cell which may explain the dysfunction induced in white adipose tissue (WAT) during obesity.

## Results

### Total FA composition and shotgun lipid profiling of adipose tissue

To place subsequent results in context, total FA compositions of the two diets fed to the mice (regular chow diet: RCD and high fat diet: HFD) were analysed by gas chromatography coupled with mass spectrometry (GC-MS) (**Fig. 1**). While the RCD contained much less fat (11.5%) than HFD (55%) in composition the RCD was dominated by C18:2 (51.6%) and C16:0 (16.4%). The HFD included a greater relative distribution of FAs, with the proportions of C18:2 and C16:0 decreasing to 34.9% and 12.8%, respectively, while other FAs, especially C18:1 (29.4%), accounted for a greater proportion than in the RCD.

**Fig 1.**
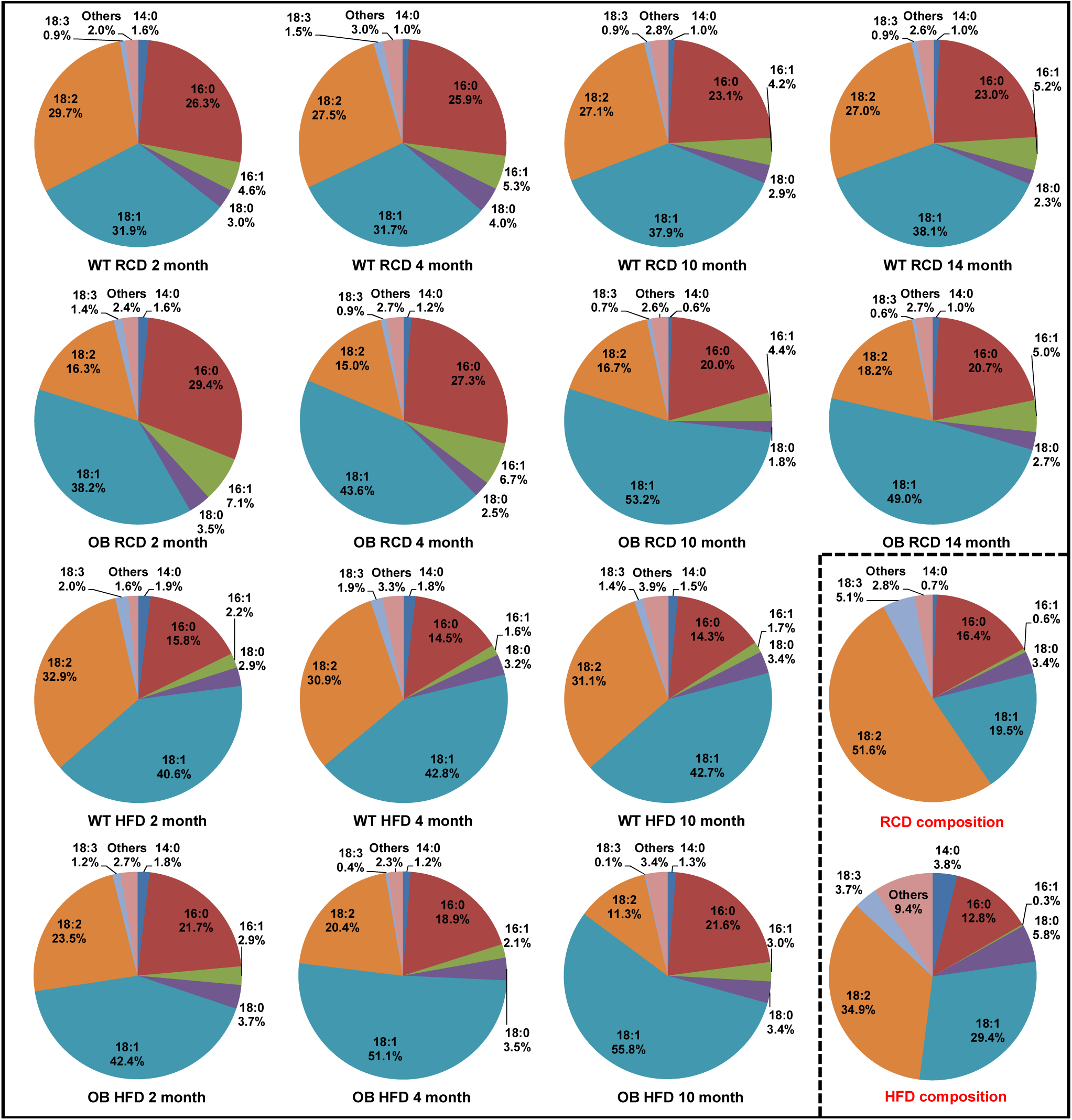
Total fatty acid composition in adipose tissues from *ob/ob* (OB) mice and wild-type (WT) controls aged between 2 and 14 months fed by regular chow diet (RCD) or a high fat diet (HFD), and total FA composition in the two diets.

These profiles were then compared with the total FA composition in WAT of all mice across the diets and time points. There were clear differences in composition between the diet and WAT from RCD-fed wildtype mice (WT-RCD) (**Fig. 1)**. The proportion of C18:2 was decreased while C16:0, C16:1 and C18:1 were increased in WAT relative to the RCD diet, indicative of increased DNL, SCD-1, and ELOVL6 activity. These changes were more marked for the total FA composition of RCD-fed *ob/ob* mice (OB-RCD), with the total fatty acid profile clearly exhibiting larger increases in C18:1 and decreases in C18:2 than the WT-RCD group. Comparing OB-RCD and WT-RCD across the time course, the relative proportion of C16:0 and C16:1 were increased in younger animals (2 and 4 month), while this proportion decreased in older animals (10 and 14 month), with C18:1 becoming more dominant.

However, for the WT-HFD mice, C16:0 and C16:1 were decreased compared with both WT-RCD and OB-RCD, whereas C18:1, C18:2, C18:3, and C14:0 were increased in all age groups of WT-HFD mice compared with the WT-RCD mice. These lipid changes largely reflected the FA composition of the HFD, although WT-HFD adipose tissues had slightly more C16:0 and C18:1 and less C18:2, presumably in part influenced by DNL of carbohydrate in the diet. In OB-HFD mice **(Fig. 1**), the FA composition was a mixture of dietary FAs, especially C18:2, and *de novo* synthesized FAs (C16:0 and C18:1). Ageing was associated with more C18:1 and less C18:2 in OB-HFD mice, presumably associated with raised DNL and SCD activity. In addition, across all four diet/mouse strain combinations C18:1 accumulated while C16:0 declined with age, suggesting more FA elongation by ELOVL6 and desaturation by SCD1 in aged mice.

To complement the total fatty acid dataset, intact lipids were determined by direct infusion (DI-) MS, with the resulting profiles dominated by neutral lipids (triglycerides (TGs), diglycerides (DGs) and sterols). The adipose tissue of *ob/ob* and WT mice fed on the RCD and HFD were readily differentiated by 2-Way Orthogonal Partial Least Squares-Discriminate Analysis (O2PLS-DA) (**Fig. 2A**, Q^2^=85%). From inspection of the scores plot, component 1 was largely associated with diet while component 2 was associated with the difference between the WT-RCD compared with the other three groups. This indicates that the HFD has a bigger influence on adipose tissue metabolism than genotype (component 1). Moreover, examining the associated loadings plots these classifications were most influenced by TG species across both the first and second components, while PLs were most associated with the first component. This suggests that while genotype and diet influence the profile of TGs, the PL composition was most influenced by diet rather than by genotype.

**Fig 2.**
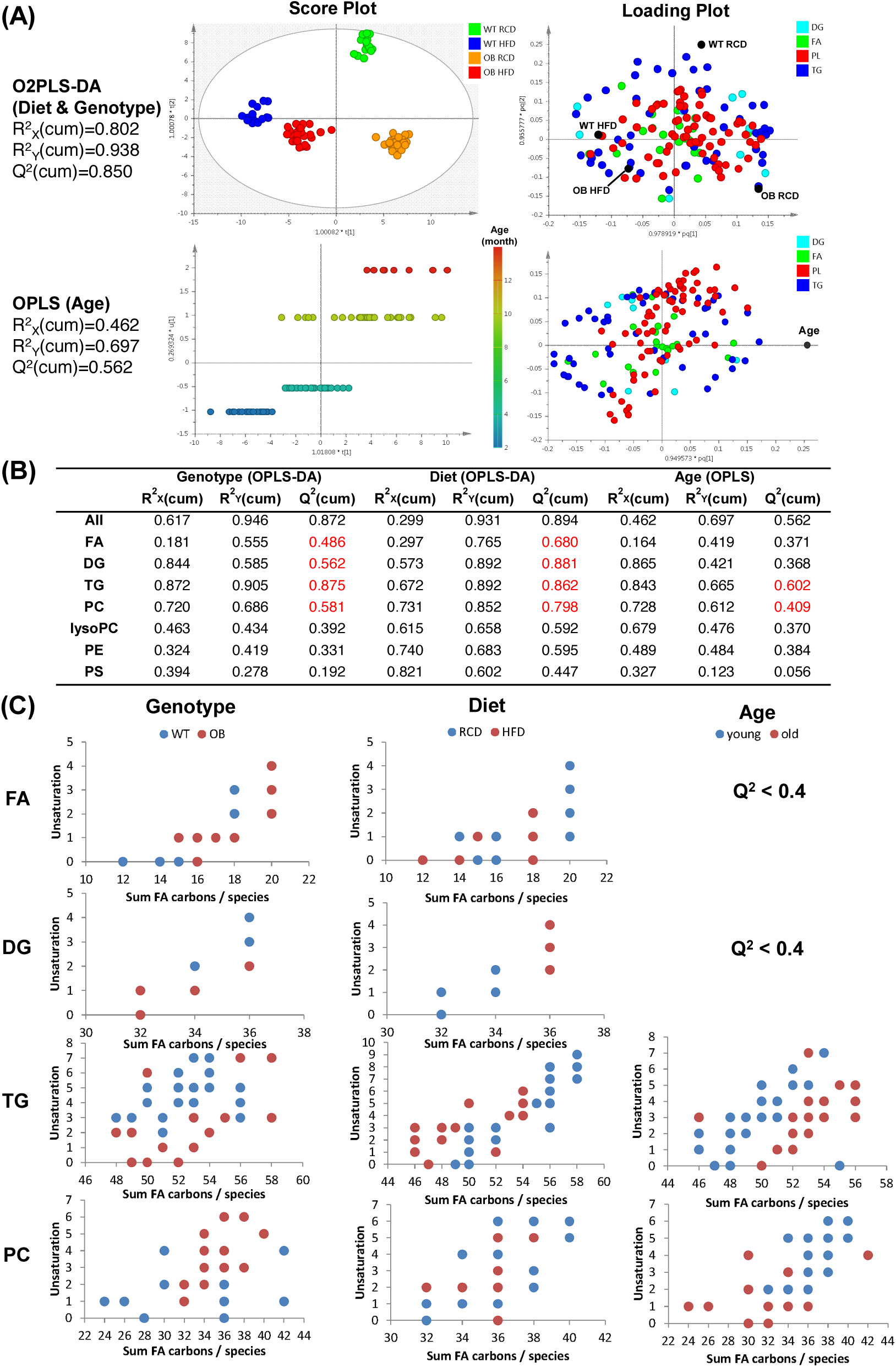
(A) Multivariate analysis of different adipose tissues by different factors (OB/WT, RCD/HFD, 2/4/10/14-month) using all classes of lipids; **(B)** OPLS-DA and OPLS model for each class of lipids in adipose tissues, and the specific lipids changed the most according to these models. **(C)** Lipid species changed most significantly by genotype/diet/age according to the models in (B).

Next, we applied Orthogonal Partial Least Squares (OPLS), a multivariate regression tool developed from the PLS algorithm, to model the lipid changes most associated with ageing and the feeding time course regardless of genotype and diet, and while this model was less robust than that for genotype and diet differences, a regression model was built (**Fig. 2A** second row, Q^2^ =56%). This regression model was largely driven by changes in TG species, reflecting the dominance of this class of lipids to the total adipose tissue lipidome.

Because neutral lipids dominate the lipidome of WAT, solid phase extraction (SPE) was performed to pre-fractionate polar lipids (lysophosphatidylcholines (lysoPCs), PCs, PEs, and phosphatidylserines (PSs)) prior to liquid chromatography (LC-) MS. In total, 157 species were measured in 7 major lipid classes (FA, DGs, TGs, lysoPC, PC, PE and PS). Next, the data collected using fractions from the SPE method was considered individually, and again OPLS-DA and OPLS models were built to evaluate the influence of the three factors (diet, genotype and age) across the study design using each of the aforementioned 7 classes of lipid (**Fig. 2B**). Neutral lipids (DG and TG) and FAs were both affected by diet and genotype as modelled by OPLS-DA (all Q^2^>48%), while for the PLs only PCs could be used to build a good model to differentiate OB/WT and HFD/RCD. In terms of modelling age, only the TG and PC subsets produced robust PLS models (Q^2^ =56% and 41%, respectively). Interrogation of the corresponding loadings plots and the variable influence on projection (VIP) for these OPLS-DA models identified individual metabolites responsible for the separation across different diets, genotypes and ages (**Fig. 2C**).

### Neutral lipids and phospholipids exhibit distinct and different changes in chain length and unsaturation in *ob/ob* mice

To further examine the lipid profiles detected across the different lipid classes, the relative proportions of lipids with the same total FA carbons or total FA unsaturation for the seven different classes were plotted (**Fig. 3** and **Fig. 4)** to provide information about how each lipid class is elongated and desaturated by genotype, diet and age interactions. The 16-carbon total FAs increased in all *ob/ob* groups compared with WT animals (**Fig. 3**), while C18 fatty acids decreased slightly. Fatty acid side chain length also decreased in the DGs and TGs of *ob/ob* mice compared with WT mice, owing to increases in 32/34-carbon DGs and 50/52-carbon TGs, and decreases in 36-carbon DGs and 54-carbon TGs in most groups (**Fig. 3**).

**Fig 3.**
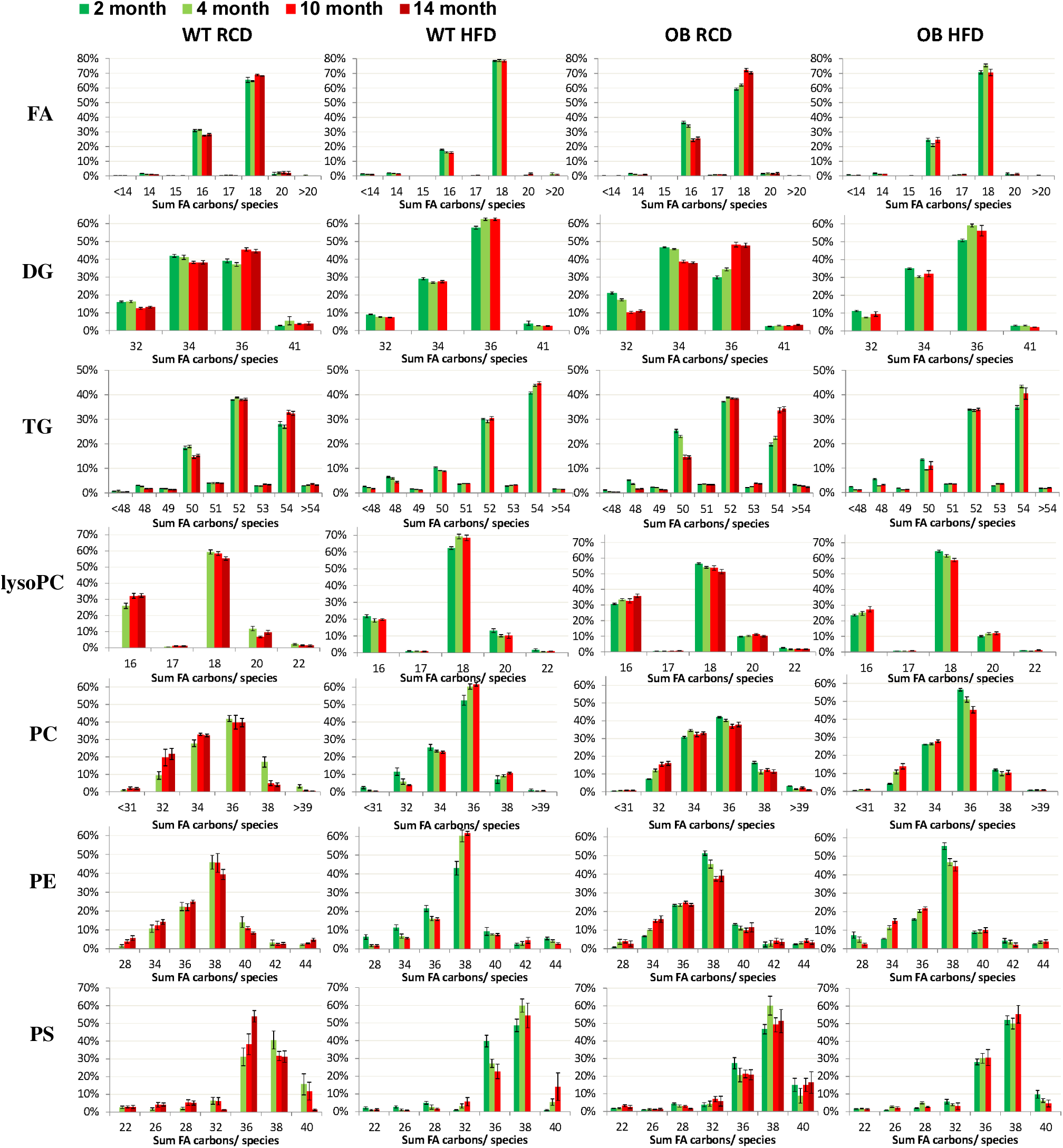
Fatty acid chain length varies in adipose tissues from *ob/ob* (OB) mice and wild-type (WT) controls aged between 2 and 14 months fed by regular chow diet (RCD) or a high fat diet (HFD).

**Fig 4.**
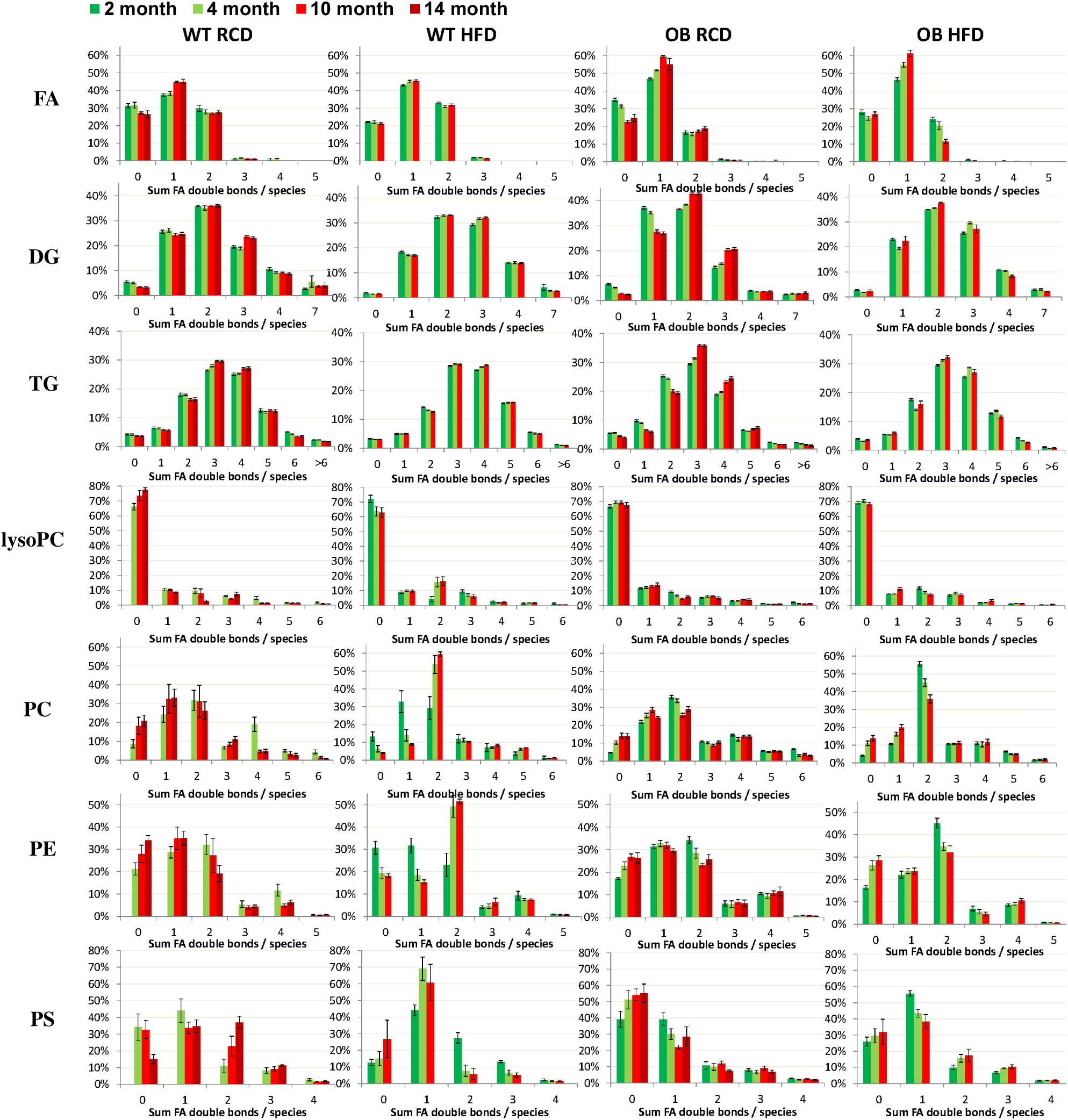
Fatty acid unsaturation varies in adipose tissues from *ob/ob* (OB) mice and wild-type (WT) controls aged between 2 and 14 months fed by regular chow diet (RCD) or a high fat diet (HFD)

Differences in the degree of unsaturation were much more striking than the changes detected in chain length. Oleic acid dominated the profiles of *ob/ob* mice (**Fig. 4**, first row), with all other unsaturated fats decreased in comparison with the WT mice, while saturated fats were on the whole increased, presumably in part by increased DNL, given that the animals were on the same diet.

While there were similarities in the differences in terms of chain length and unsaturation between WT and *ob/ob* mice for the classes of FA, DG and TGs, PLs, especially PCs, exhibited very different changes between WT and *ob/ob* mice. The PC fraction consisted of fatty acids with slightly longer and much more unsaturated chains in *ob/ob* mice compared with WT. One of the most marked changes was an increase in PCs where the degree of unsaturation was more than two double bonds (DU>2) in *ob/ob* mice on a RCD compared with WT, and PCs with DU=2 for both strains on a HFD (**Fig. 4**). Comparing profiles between the TG and PC classes it was striking that despite the increase in C16:0 and C18:1 incorporated into neutral lipids and FAs, more highly unsaturated FAs (DU>2) were selectively incorporated into PCs. Furthermore, more long-chain FAs (>18 carbons) were incorporated into PEs and PSs (**Fig. 3**) compared with other lipid classes (FA, DG, TG, lysoPC, PC).

### HFD is the determining factor for lipid composition in mouse WAT

When considering the chain lengths of TG and DG species following a HFD, both WAT from WT and *ob/ob* mice were dominated by species containing more 18-carbon FA side chains and less 16-carbon FA side chains (**Fig. 3**), particularly the accumulation of 54-carbon TGs (mostly comprised of three 18-carbon FA chains) at the expense of 50-carbon TGs (mostly composed of two 16-carbon FAs and one 18-carbon FA) and 52-carbon TGs (one 16-carbon and two 18-carbon FA chains). A similar pattern was also detected for 32, 34, and 36-carbon DGs. For neutral lipids there was only a modest increase in unsaturation associated with HFD (**Fig. 4**).

For PLs, the chain length changes of lysoPCs and PCs induced by a HFD showed a reduction in the distribution of lipids, with 18-carbon lysoPCs and 36-carbon PCs dominating the profiles compared with the RCD fed animals (**Fig. 3**), along with a marked reduction in 20-carbon FA containing species. There was also an overall increase in unsaturated fatty acid side chains for both lysoPCs and PCs (**Fig. 4**).

### Longer TGs and saturated PCs accumulate in WAT of aged mice

As detected by GC-MS (**Fig. 1)**, lipids containing C18:1 fatty acids increased while C16:0 decreased with age. These ageing trends are mostly reflected in the accumulation of 54-carbon TGs and a decrease in 50-carbon TGs in older mice, along with similar trends in DGs (**Fig. 2C** & **Fig. 3**). However, PCs demonstrated a very different profile of change with age compared with neutral lipids. With age PCs containing either saturated or monounsaturated fatty acids accumulated in WAT of older mice.

### Liquid chromatography-ion mobility-mass spectrometry provides increased resolution of lipid species and confirms major lipid trends

While the SPE strategy improved the detection of a larger number of lipid species from the adipose tissue extracts compared with direct measurement of the total lipid extract, many lipids were either isocratically eluted, isobaric or demonstrated a combination of the two properties. Thus, the LC-MS dataset collected on all the adipose tissue samples underestimates the true complexity of the data. To address this, liquid chromatography-ion mobility-mass spectrometry (LC-IM-MS) was performed on the SPE fractions collected from adipose tissue from 10 month old animals. As with the LC-MS dataset, the LC-IM-MS dataset demonstrated that the free FA and PL fractions were largely dependent on the genotype of the animals, while the neutral lipid fraction was determined by both genotype and diet (**Supplementary Figure 1A**). In the phospholipid fraction a total of 71 features were significantly changed with a p<0.05 and fold change > 2 (**Supplementary Figure 1B**) and were annotated according to exact mass match with the LIPID MAPS Structure Library (LMSD) (23). As classes of lipids, unsaturated PLs and both unsaturated and saturated LPLs increased in *ob/ob* mice, regardless of diet (**Supplementary Figure 1C**). In the neutral lipid fraction, 57 lipids classified as triglycerides, diglycerides and ceramides were significantly changed with a p<0.05 and fold change > 2 (**Supplementary Figure 1B**) and were annotated using the LMSD. Unlike the PL fraction, neutral lipids were altered by both genotype and diet (**Supplementary Figure 1B**).

### One-carbon metabolism in mice changes significantly with genotype and correlates with changes in phosphatidylcholine metabolism but not triglycerides

Metabolites involved in the 1-carbon cycle were measured in WAT by LC-tandem MS (LC-MS/MS). Differences between genotype and diet became more pronounced in older mice, especially at 10-months (**Fig. 5A)**. All WT-HFD mice contained fewer 1-carbon metabolites in their adipose tissue than WT-RCD, while for *ob/ob* mice this pattern was less clear.

**Fig 5.**
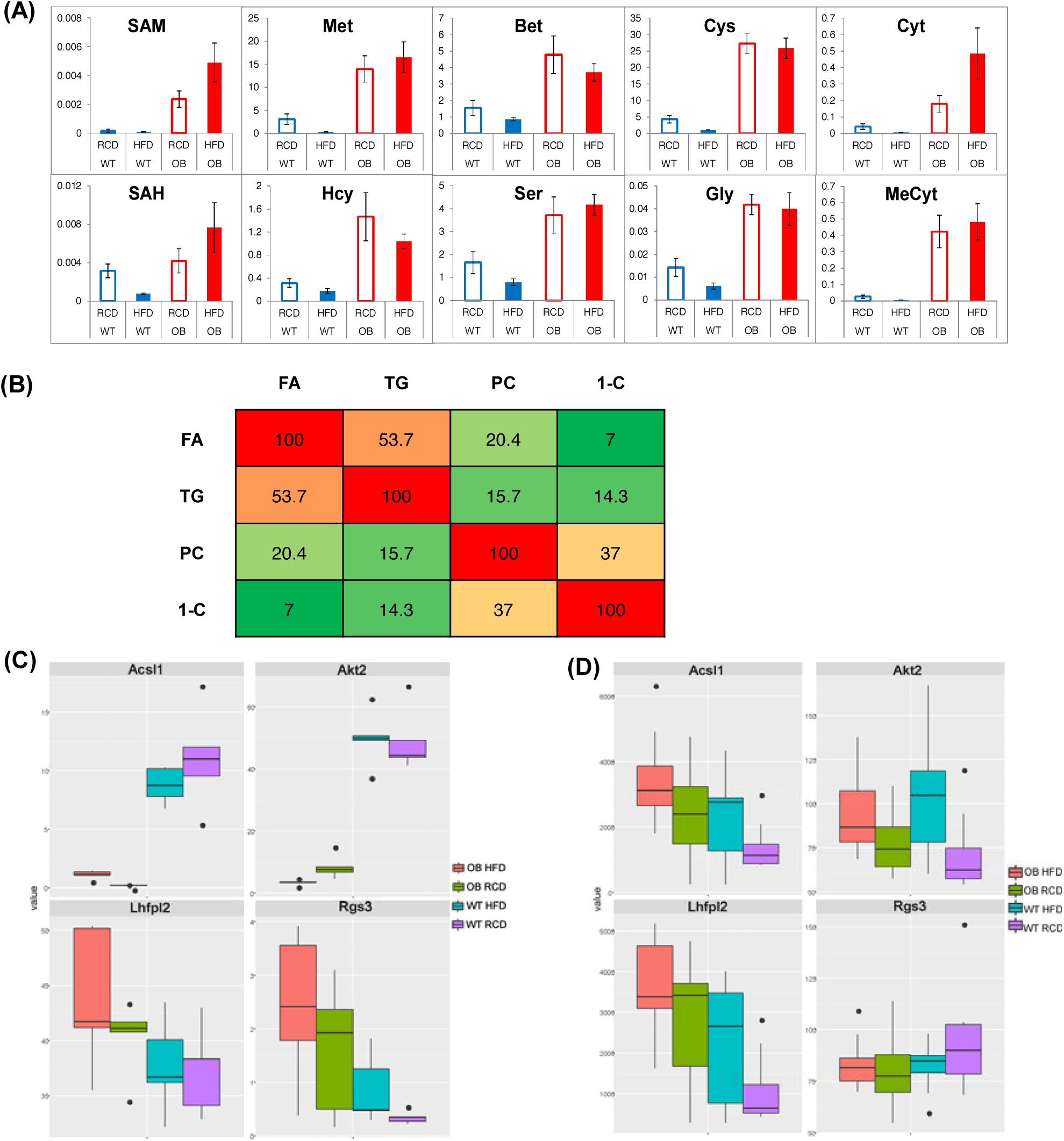
(A) 1-Carbon metabolites varied in adipose tissues from 10-month *ob/ob* (OB) mice and wild-type (WT) controls fed a regular chow diet (RCD) or high fat diet (HFD); **(B)** Correlation (Q^2^_cum_, %) of different metabolic data by O2PLS application; **(C)** DNA methylation and **(D)** expression of four genes selected from literature.

Given that choline is a central metabolite in the 1-carbon cycle, we hypothesized that lipid metabolism might correlate with changes in the 1-carbon cycle. OPLS was used for data integration combining different metabolite data. With this methodology, systematic variation that overlaps across analytical platforms can be separated from platform-specific systematic variation to validate the correlation between the datasets. FAs, TGs, PCs and 1-C metabolites were studied using this method to understand changes occurring as a result of genotype, diet and age. Total FA data (as measured by GC-MS) and TG data (as measured by DI-MS) were correlated well (Q^2^=53%), as would be expected as the total FA profile detected by GC-MS will be dominated by fatty acids in TGs (**Fig. 5B**). Of more note, the 1-C cycle metabolite data correlated better with PCs (Q^2^=37%) than with total FAs (Q^2^=7%) and TG (Q^2^=14%). This suggests that when PC metabolism is altered during obesity this has a consequential effect on 1-C metabolism, potentially changing the supply of SAM for processes such as DNA methylation. The conversion between PCs and PEs also relies on SAM. Furthermore, the correlations between 1-carbon cycle and PCs were greater in *ob/ob* mice than in wild type mice (data not shown).

### Leptin deficiency results in significant changes in DNA methylation status

As SAM provides methyl groups for DNA methylation, we also determined the methylation status of four genes which are functionally related to obesity and type 2 diabetes and known to be partly under epigenetic control – Acyl-CoA Synthetase Long Chain Family Member 1 (ACSL1), serine/threonine-protein kinase 2 (AKT2), Regulator Of G Protein Signaling 3 (Rgs3) and Lipoma HMGIC fusion partner-like 2 (Lhfpl2) (24-26). ACSL1 and AKT2 were much less methylated in *ob/ob* mice, while Rgs3 and Lhfpl2 were more methylated (**Fig. 5C**) compared with WT. In *ob/ob* mice, from our LC-MS/MS analysis, there was an increase in the pool of SAM for DNA methylation which may contribute to the alterations in methylation status of the genome between WT and *ob/ob* mice. Interestingly, despite the large differences in choline content between the RCD and HFD, the HFD had a smaller impact on 1-carbon metabolites and the methylation status of the four genes examined when compared with genotype changes.

Expression of these four genes were also determined (**Fig. 5D**) and again genotype had a major effect. There was increased expression of Acsl1 alongside reduced methylation in *ob/ob* mice, but expression of Lhfpl2 was also increased despite being more methylated in *ob/ob* mice. However, the expression of Rgs3 and Akt2 were not significantly changed by genotype. Moreover, the influence of the HFD on gene expression was much greater than its effect on the methylation status, with all four genes exhibiting increased expression in HFD-fed mice.

Finally, to examine global methylation, DNA was extracted from adipose tissue from 10 month old animals (WT and *ob/ob*, RCT and HFD), digested and the ratio of methyl-cytosine to cytosine calculated using LC-MS/MS (WT-RCD = 2.5%, *ob/ob*-RCD = 4.0%, WT-HFD = 3.1%, *ob/ob*-HFD = 6.1%). Two-way ANOVA demonstrated that genotype was associated with a significant change (p=0.001), but not diet nor the interaction between genotype and diet.

### Transcriptomics analysis

To better understand what molecular changes were associated with the lipidome changes detected in the two mouse strains and by the two diets, a transcriptomic study was conducted to compare the differences in gene expression of adipose tissues. Raw data were subject to quantile normalisation with the detection P-value threshold set to 0.01. Gene expression was compared with 95% (p<0.05) and 99% (p<0.01) confidence intervals for each factor (genotype, diet and age), as shown in **Table 1**, with 5341 genes found to be significantly changed.

**Table I.** Transcriptomic summary of gene expression changed significantly (p<0.05/p<0.01) in adipose tissues from *ob/ob* (OB) mice and wild-type (WT) controls aged between 2 and 10 months fed by regular chow diet (RCD) and high fat diet (HFD).

| Comparison | Group | Number of selected probes | Significant results at 0.05 | Significant results at 0.01 |
| --- | --- | --- | --- | --- |
| **WT vs. OB** | RCD 2 month | 15596 | 2511 | 121 |
|  | RCD 10 month | 15392 | 2309 | 1122 |
|  | HFD 2 month | 16413 | 1303 | 465 |
|  | HFD 10 month | 15245 | 2539 | 1172 |
| **RCD vs. HFD** | WT 2 month | 16572 | 2 | 0 |
|  | WT 10 month | 14902 | 878 | 19 |
|  | OB 2 month | 15551 | 361 | 37 |
|  | OB 10 month | 15566 | 1 | 0 |
| **2 month vs. 10 month** | WT RCD | 15347 | 5 | 2 |
|  | WT HFD | 16305 | 84 | 4 |
|  | OB RCD | 15308 | 1264 | 456 |
|  | OB HFD | 15346 | 1722 | 726 |

Genotype resulted in the greatest number of significantly changed genes (2077 transcripts for the RCD at 2 months, 1086 transcripts for HFD at 2 months, 1895 transcripts for RCD at 10 months, and 2063 transcripts for HFD at 10 months), while HFD had a more modest impact on gene expression (2 transcripts for WT at 2 months, 311 transcripts for *ob/ob* at 2 months, 761 transcripts for WT at 10 months, and 1 transcript for *ob/ob* at 10 months). The relative increase in obesity might be a reason for this discrepancy with genotype causing a larger change than diet in the mouse study. Ageing also resulted in more transcriptional changes in *ob/ob* mice compared with controls (5 transcripts for WT on a RCD, 80 transcripts for WT on a HFD, 1079 transcripts for *ob/ob* on a RCD, and 1428 transcripts for *ob/ob* on a HFD).

To further explore the specific pathways regulated by the *ob/ob* genotype, HFD diet, and aging, KEGG Pathway Enrichment analysis was conducted. The inflammatory and immune system pathways dominated the up-regulated terms (**Supplementary Tables 1 and 2**). Inflammation-related pathways were significantly up-regulated by the *ob/ob* genotype in all but the HFD-10 group, by the HFD diet in the WT-10 group and by aging in the OB-RCD group. Inflammation wasn’t associated with all of the *ob/ob* and HFD comparisons, perhaps suggesting that there may be an upper level to the effect of inflammation at the transcriptional level.

The *ob/ob* genotype was associated with the lysosome, phagosome, and signaling pathways that play a key role in the immune system, such as toll-like receptor, nuclear factor kappa-light-chain-enhancer of activated B cells (NF-κB) and chemokine signaling pathways. The KEGG pathways enriched by genes significantly up- or down-regulated by HFD diet and ageing are also similar to those enriched by *ob/ob* genotype, in particular inflammatory pathways and remodeling associated with organelles like the phagosome and lysosome pathways. Ageing was also associated with down-regulation of peroxisome pathways and metabolic pathways in general, including carbon metabolism and fatty acid metabolism.

To further examine the data, we selected some genes known to be involved in the pathways of DNL (FASN, desaturation (SCD-1), endoplasmic reticulum stress (XBP1, EDEM1, and CARMIL3), adipose tissue remodelling (AGPAT6, MBOAT1, MBOAT2) and related transcription factors (INSIG1, SREBP-1c, SREBP-2, PPARα, PPARγ, PPARδ, and MYC) to determine variations in expression (**Fig. 6**). The significance of these expression changes is shown in **Fig. 7A**. Fasn and Edem1 increased in expression in *ob/ob* mice and HFD-fed mice. Expression of remodelling genes (Agpat6, Mboat1, and Mboat2) decreased in *ob/ob* and HFD-fed and aged mice, suggesting that WAT remodelling is impaired by both obesity and ageing. Other significant changes included decreased expression of PPARα in *ob/ob* mice and decreased expression of Srebp2 in HFD-fed mice, representing different pathways altered by genotype/diet-induced obesity. Moreover, we examined how these genes are correlated with each other in terms of expression changes (**Fig. 7B**). Adipose tissue remodelling genes (Mboat1, Mboat2) correlated strongly with Srebp2. Lipid metabolism related genes (Fasn, Scd1, Xbp1 and Edem1) correlated with transcription factors Srebp1, PPARα, PPARγ, PPARδ, and Myc.

**Fig 6.**
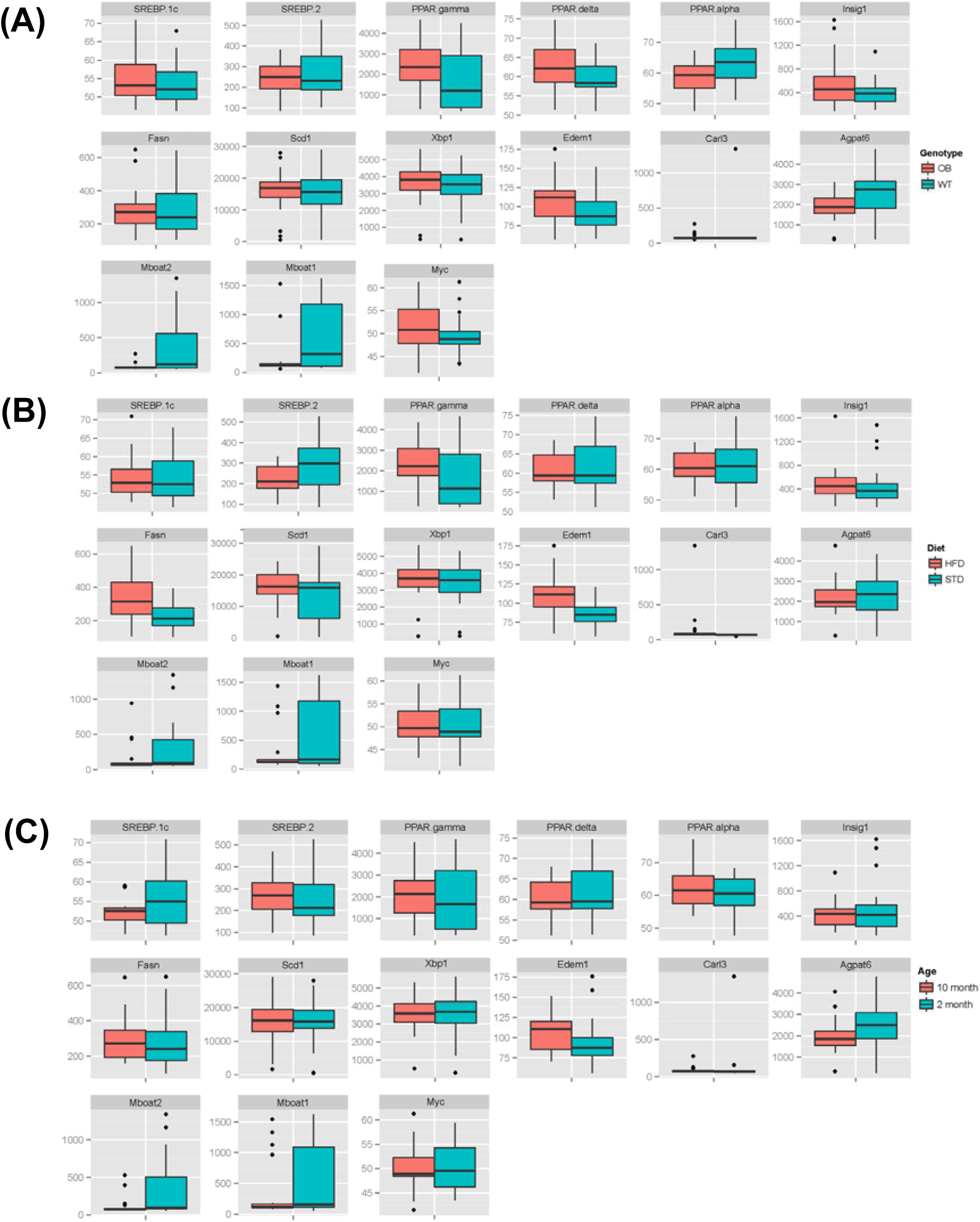
(A) Selected gene expression comparison between *ob/ob* (OB) and wildtype (WT) mice; **(B)** Selected gene expression comparison between high fat diet (HFD) and regular chow diet (RCD) fed mice; **(C)** Selected gene expression comparison between 2 month and 10 month old mice.

**Fig 7.**
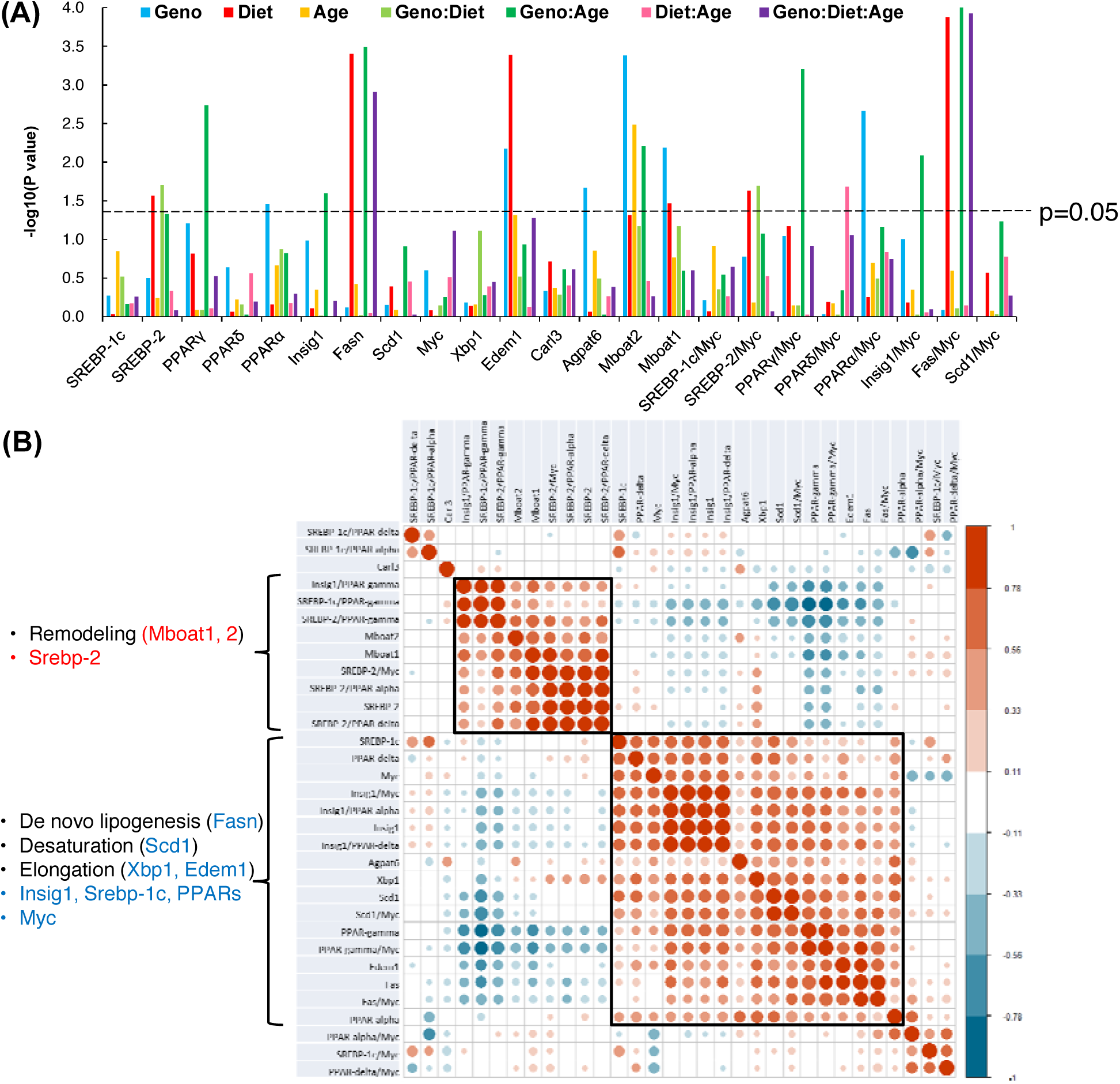
(A) Significance (P-value) of selected gene expression comparison in terms of genotype/diet/age groups factors; **(B)** Heatmap showing correlation of selected genes involved in de novo lipogenesis (DNL), desaturation, elongation, adipose tissue remodelling and related transcription factors.

The transcriptomic patterns of Fasn, Insig1, Insig2, PPARγ, and Srebf1 were highly similar, and up-regulated in adipose tissue of *ob/ob* mice at the age of 2 months compared with wild-types but down-regulated at 10 months compared with wild-type controls, up-regulated in adipose tissue of 10 month old wild-type mice during aging but down-regulated in 10 month old *ob/ob* mice compared with the younger animals. This suggests that DNL was activated by leptin deficiency in younger mice and with aging in wild-type animals, but inhibited in adipose tissue of old *ob/ob* mice. Scd1 and Elovl6 were regulated in similar directions with Fasn which may drive the lipidome remodelling detected.

Expression of Elovl1 was up-regulated by *ob/ob*, the HFD and aging, while Elovl6 was down-regulated significantly by HFD in the same groups, suggesting elongation of C18:0 by ELOVL1 is activated to form more long chain fatty acids in adipose tissue of either old or obese mice. However, further desaturation of oleate was limited as the desaturase genes (Fads1 and Fads2) were down-regulated by *ob/ob*, HFD, and aging. This agrees with the dominance of oleate in adipose tissue of obese and aged mice. Moreover, the expression of TG biosynthesis and lipid remodelling genes (Mboat1 and Mboat2) were down-regulated by obesity from either *ob/ob* or HFD.

## Discussion

Stored lipids in adipose tissue, largely TGs, are either derived from the diet, or associated with DNL and subsequent modifications. Here, both mouse strains on a RCD diet (11.5% fat) demonstrated DNL activity as shown by the TG content of their WAT, with the dominant components of adipose TGs being C16:0, C16:1, and C18:1, despite C18:2 being the major fatty acid in the RCD. This was even more apparent in *ob/ob* mice, where leptin deficiency increases appetite and led to an increased production of C18:1 oleic acid, showing concerted action on ACC, FAS, SCD-1 and ELOV6, which have a higher flux than those enzymes involved in subsequent fatty acid modifications. The activities of Δ^5^-eicosatrienoyl-CoA desaturase and Δ^6^-oleoyl(linolenoyl)-CoA desaturase are reported to be associated with hyperglycemia and non-alcoholic steatohepatitis, and palmitate can compete with omega-6 linoleic and omega-3 α-linolenic acids for FADS2 mediated Δ^6^-desaturation (27-29).

There are conflicting reports as to whether an increased oleate-content of WAT is protective or deleterious to whole body insulin resistance (30, 31) (32). We have reported previously (33) that TGs containing fatty acids with 16-18-carbons and derived from Δ9 desaturation, especially oleate, were increased in *ob/ob* mice. In this previous study, while AdTG-*ob/ob* mice had increased SCD ratios but decreased Elovl6 ratios in WAT, the mice were more obese but relatively metabolically healthy compared with *ob/ob* mice, while the AKT2 KO mice which are lean but insulin resistant had increased Elovl6 ratios but not significantly changed SCD ratios compared with wildtype mice. This suggests that oleic acid production may be protective for insulin resistance, but at some stage this is overwhelmed by increasing lipid storage, possibly related to the decline in proportion of essential n-3 and n-6 PUFAs.

While relatively increased fatty acids C16:0 and C18:1, and decreased C18:2 fatty acid content was also apparent in DGs and FAs in *ob/ob* WAT, this pattern did not extend to PCs which had longer and more unsaturated fatty acids, particularly highly unsaturated fatty acids (HUFAs). HUFAs, in contrast to MUFAs and precursor PUFAs, are usually found in membrane lipids as components of PLs where they contribute to maintenance of membrane fluidity and sensitivity to hormones, and also regulate gene expression through various transcription factors, such as the PPAR receptors, liver-X receptor, HNF4, and SREBP-1c (34-36). The selective enrichment of PLs with HUFAs could be another protective mechanism to maintain normal function and membrane fluidity of adipocytes during large consumption of carbohydrates which in turn activates DNL.

WAT from WT-HFD mice reflected the FA composition of the HFD, except for a moderate increase in C16:0 and C18:1, presumably due to a contribution from DNL, particularly for FAs, DGs and TGs. The pattern was also apparent for lysoPCs, PCs, PEs, and PSs, and this contributed to a decrease in the distribution of lipids in the HFD-fed mice due to domination of a relatively small number of dietary FA species. While the WT-HFD group appeared to ‘resist’ the adoption of the dietary fatty acid profile by maintaining a greater range of lipids, the OB-HFD group had the greatest reduction in lipid species compared with the other groups. This loss of diversity may have a major impact on cellular function, particularly of cell membranes. Previously we have shown (37) that myocardial mitochondrial dysfunction was detectable in the hearts of OB-HFD mice (3 month) much earlier than in OB-RCD mice (14 month) which may reflect changes in sub-cellular membrane composition.

One of the most profound changes in metabolism detected was the increase in 1-C cycle metabolites, and in particular SAM for methylation, in *ob/ob* mice on a RCD compared with wildtype mice. While a HFD decreased all 1-C metabolites in wildtype mice it produced a more complex pattern in *ob/ob* mice with less homocysteine, betaine, cysteine, and glycine but more SAM, SAH, methionine, serine, cytosine, and methyl-cytosine than OB-RCD mice. The CDP-choline pathway links choline to phosphocholine formation as well as the 1-C pathway (38). SAM can also be used to methylate PE to form PC. Moreover, SREBP-1, which is induced by insulin signalling, can interact with SAM through 1-C metabolism and lipid metabolism (19, 39, 40). These pathways suggest crosstalk between lipid metabolism and SAM formation. In obese mice, excess fat storage results in a greater demand for PCs to be utilized to form and enlarge membranes to support adipocyte expansion and differentiation. In *ob/ob* mice the whole choline/1-carbon metabolism pathway may be activated to support PC formation, in part to drive cell membrane formation. This upregulation in SAM production may also alter the methylation capability for DNA and histones, explaining some of the methylation changes we detected in key transcription factors, ACSL1, AKT2, Rgs3, and Lhfpl2, whose epigenetic changes in WAT are known to alter systemic metabolism (24-26).

Transcriptomic changes demonstrate that a number of lipid regulating and remodelling genes were repressed by leptin deficiency, HFD, and aging including decreased expression of PPARα in *ob/ob* mice and decreased expression of Srebp2 in HFD-fed mice. Coordinated remodelling of the structure and metabolism of WAT has been reported to play an important role in developing insulin resistance and cardiovascular diseases (41-43). Using correlation analysis, the remodelling genes correlate well with Srebp2 while lipid synthetic genes were correlated with Srebp1c and PPARs in the current study. Srebp2 regulates cholesterol synthesis and sterol content of cell membranes, while Srebp1c regulates fatty acid synthesis (44). Carobbio and colleagues (15) showed that Insig1 mRNA expression decreased in WAT from mice with obesity-associated insulin resistance and in morbidly obese humans. Insig1 downregulation promotes the maintenance of mature SREBP1 and facilitates DNL and FA unsaturation, compensating for the anti-lipogenic effect of insulin resistance. In addition all PPARs play central roles in regulating carbohydrate and lipid metabolism in WAT and the decrease in PPARα and increase in PPARγ is consistent with a switch from fatty acid oxidation to storage and elongation in WAT during increased storage of TGs (45).

In the present study, we selected important variables from lipidome, metabolome and one carbon metabolism using VIP scores. By selecting these variables using VIP scores, we reduce the contribution of noise and variables with high variation, and this step is essential for translational research to target candidate variables from large scale -omics data sets (46).

In summary, we demonstrated that on a RCD both wildtype and *ob/ob* mice regulate fatty acid composition differently between storage lipids, largely TGs and DGs, and the phospholipid components of the cell membrane, presumably reflecting the importance of maintaining cell membrane function. Conversely, most of the lipids stored in the WAT of HFD-fed mice are assembled directly using dietary fatty acids. This has far-reaching effects including alterations in 1-C metabolism, epigenetic changes and transcriptionally including key transcription factors involved in coordinating lipid metabolism.

## Materials and methods

### Animals and diets

Five-week old male *ob/ob* mice and their wild type (C57B1/6J, WT) controls were purchased from a commercial breeder (Harlan UK). Mice were housed in a temperature– and humidity-controlled facility, with a 12-h: 12-h light-dark cycle and access to water ad libitum. Mice (n=10/age group/genotype) fed on regular chow diet (Caloric content: 11.5% fat, 26.9% protein, 61.6% carbohydrate) (RM1, Special Diet Services, Witham, Essex, UK) were sacrificed at the age of 2-months, 4-months, 10-months and 14-months. A separate group of mice were switched to a custom-produced high fat diet (Caloric content: 55% fat, 29% protein, 16% carbohydrate; fatty acid composition: 27% saturated fatty acid, 48% mono unsaturated fatty acid and 25% poly unsaturated fatty acid) (diet code: 829197; Special Diet Services, Witham, Essex, UK) at different stages for various lengths of time and were sacrificed at the age of 2-months (3-week high fat feeding) 4-months (12-week high fat feeding) and 10 months (12-week high fat feeding). This diet has been previously shown to be obesogenic in mice (47). Mice were euthanised by carbon dioxide asphyxiation, and upon cessation of peripheral signs WAT was rapidly dissected, snap frozen with liquid nitrogen, and stored at −80°C until further analysis. All animal procedures were performed under a project license and approved by the UK Home Office and University of Cambridge Animal Welfare Ethical Review Board.

### Sample Pre-treatment

#### FAs, TGs and DGs

Metabolites were extracted from tissue using a modified Folch method (48). In brief, ∼20 mg frozen tissue was pulverised in chloroform-methanol (400 µl; 2:1 v/v) using a TissueLyser (Qiagen), then the mixture was sonicated for 10 minutes. Water (80 µl) was added and samples were then centrifuged (13,200 rpm, 10 min). The resulting aqueous and organic layers were collected in separate glass tubes. The pellet was extracted two further times and the fractions pooled. Organic extracts were dried overnight under nitrogen gas, and aqueous extracts were evaporated to dryness using an evacuated centrifuge (Eppendorf, Hamburg, Germany).

#### PLs and 1-carbon metabolites

Intact lipids were extracted by a modified Bligh and Dyer (BD) method (49) from WAT (∼50 mg frozen tissue) to separate aqueous-soluble metabolites from lipids. Approximately ∼50 mg frozen tissue was pulverised in methanol-chloroform (300 µl, 2:1 v/v) using a TissueLyser (Qiagen, West Sussex, UK). Then the mixture was sonicated for 10 min. Chloroform-water (1:1) was added (100 µl of each component) and samples were then centrifuged (16,600 rcf, 20 min). The resulting aqueous and organic layers were collected in separate tubes. This procedure was repeated twice. Organic extracts were dried overnight under nitrogen gas, and aqueous extracts were evaporated to dryness using an evacuated centrifuge (Eppendorf, Hamburg, Germany).

An SPE method was used to separate different fractions of lipids from organic extracts. CHROMABOND® NH_2_ (3 ml, 500 mg) columns were washed with 10 ml methanol and then conditioned using 10 ml hexane before sample application. The dried organic extract was reconstituted in 1 ml n-hexane, and slowly aspirated through the column. The fraction of cholesteryl esters was firstly eluted with 1 ml hexane. Then all TGs were eluted from the column using 2 ml hexane – ethyl acetate (85:15, v/v). Finally, the column was continuously eluted using 1 ml chloroform-methanol (2:1, v/v) and 2 ml methanol to obtain the fractions of monoacylglycerides and PLs. Each fraction was dried under a flow of nitrogen in a fume hood.

### GC-MS analysis for total Fatty Acids

Organic-phase metabolites were derivatized by acid-catalysed esterification [140]. Chloroform-methanol (1:1 v/v; 400 µl) and BF_3_–methanol (10%; 125 µl) (Sigma-Aldrich, Dorset, UK) were added to the dried organic phase and incubated at 80 °C for 90 min. Water (500 µl; milliQ) and hexane (1000 µl) were added and samples were vortex-mixed for 1 min. The organic layer was evaporated to dryness in a fume hood, then reconstituted in hexane (1000 µl) for GC-MS analysis. GC-MS analyses were made using a Trace GC Ultra coupled to a DSQ2 single-quadrupole mass spectrometer (ThermoScientific). The derivatized organic samples were injected with a split ratio of 50 onto a 30 m x 0.25 mm x 0.25 µm TR-FAME stationary phase column (ThermoScientific, Hemel Hampstead, UK). The injector temperature was set to 230 °C and helium carrier gas was at a flow rate of 1.2 ml/min. The initial column temperature was 60 °C for 2 min, increased by 15 °C /min to 150 °C and then increased at a rate of 4 °C /min to 230 °C (transfer line =240 °C; ion source=250 °C; electron ionisation =70 eV). The detector was turned on after 240 s, and full-scan spectra were collected using three scans/s over a range of 50 to 650 *m/z*.

GC-MS chromatograms were processed using Xcalibur (version 2.1; ThermoScientific). Each individual peak was integrated and then normalized. For GC-MS data, peak assignment was based on mass fragmentation patterns matched to the National Institute of Standards and Technology database of mass spectra. Identification of metabolites from organic phase GC-MS analysis was supported by comparison with a FAME standard mix (Supelco 37 Component FAME Mix; Sigma Aldrich).

### UPLC-MS analysis for phospholipids

Chromatography was performed on an ACQUITY UPLC System (Waters Corporation, Elstree, Hertfordshire, UK) equipped with an Acquity UPLC 1.8 μm HSS T3 column (2.1 × 100 mm; Waters Corporation) coupled to a Waters Q-Tof Xevo mass spectrometer (Waters MS Technologies, Ltd., Manchester, UK). Column temperature was kept at 55°C.

For analysis of TGs and total extracts, the desolvation gas temperature was 500 °C, the capillary voltage was 3 kV and the cone voltage was 50 V. The binary solvent consisted of solvent A (10 mM ammonium acetate, 0.1% formic acid) and solvent B (analytical grade acetonitrile/isopropanol (1:9), 10 mM ammonium acetate, 0.1% formic acid). The temperature of the sample organiser was set at 4°C. Mass spectrometric data was collected in centroid mode over the mass range of *m/z* 50-1200 with a scan duration of 0.2 s. As lockmass, a solution of 2ng/µl (50:50 acetonitrile: water) leucine enkephaline (*m/z* 556.2771) was infused into instrument at 3 µl/min. Dried organic phase samples were reconstituted in methanol-chloroform (1:1100 µl), then diluted 20-fold further prior to injection onto the column. The gradient started from 60% A/40%B, reached 100% B in 10 min, went back to the initial conditions in 0.1 min and remained there for the following 2 min. The eluent flow rate was 0.400 ml/min.

The phospholipids extracts (fraction D from the SPE) were reconstituted in 200 µl initial mobile phase (isopropanol/acetonitrile/water, 2:1:1) prior to injection of 3 µl into the column. A lyso-phosphatidylcholine C17:0 internal standard (in initial mobile phase) was spiked into each sample to give a final concentration of 20 μM. The binary solvent system used was solvent A (HPLC-grade acetonitrile: water 60:40, 10 mM ammonium formate) and solvent B (HPLC-grade acetonitrile: isopropanol 10:90, 10 mM ammonium formate). The gradient: 0 min, 40% B, 2 min, 43% B, 2.1 min, 50% B, 12 min 54% B, 12.1 min, 70% B, 18 min, 99% B. The eluent flow rate was 0.400 ml/min. The temperature of the sample organizer was set at 4°C. The electrospray source was operated in positive ion mode with the source temperature set at 120°C and a cone gas flow of 50 L/h. The desolvation gas temperature was 550°C and the nebuliser gas flow rate was set at 900 L/h. The capillary voltage was 2 kV and the cone voltage was 30 V. Mass spectrometry data was collected in centroid mode over the mass range of *m/z* 200–1200 with scan duration of 0.2 s. As lockmass, a solution of 2 ng/μl (50:50 acetonitrile: water) leucine enkephaline (*m/z* 556.2771) was infused into the instrument at 3 μl/min.

Tandem mass spectrometry was performed as an additional function to allow fragmentation data to be collected. The MS/MS method was operated with collision energy ramp starting at 20 eV and finishing at 40 eV. The data dependent acquisition method was performed on a pool sample, in which the most abundant 5 peaks from a survey scan were selected, and the corresponding ions were subjected to MS/MS analysis.

UPLC-MS data were processed using Micromass Markerlynx Applications Manager (Waters Corporation). Peaks were detected and then matched, and retention time aligned across the samples. Lipid species were identified using exact mass, tandem mass spectrometry data and data dependent acquisition (DDA) data.

### Liquid chromatography ion mobility mass spectrometry

SPE lipid fractions were reconstituted in chloroform/methanol (1:1, v:v) and diluted with isopropanol/acetonitrile/water (2:1:1, v:v:v) solution before being analysed on an Infinity II UHPLC coupled to an Agilent 6560 Ion Mobility Q-ToF LC-MS (Agilent, Santa Clara, USA), using a 20 min reverse-phase chromatographic separation with a C18 Acquity column (1.7 μm, 2.1 × 100 mm, Waters Ltd, UK) held at 55 °C during analysis. Chromatographic separation was achieved with a solvent gradient starting from 60% solvent A (water/acetonitrile 40:60, 10 mM ammonium formate in positive ion mode, 10 mM ammonium acetate in negative ion mode) and 40% solvent B (isopropanol/acetonitrile 90:10, 10 mM ammonium formate in positive ion mode, 10 mM ammonium acetate in negative ion mode), with a flow rate of 400 µL/min. Briefly, solvent A decreases from 60% to 1% in 18 min, back to 60% in 0.5 min, followed by 2 min equilibration time.

The Dual Agilent Jet electrospray ionization source was operated at a gas temperature of 325 °C, drying gas of 5 L/min, sheath gas temperature of 275 °C, and a sheath gas flow of 12 L/min.

Drift tube and trap funnel pressure were stable at 3.9 and 3.8 Torr, respectively, while drift tube temperature was found to be stable at 23.5 °C across all acquisition runs.

Data was processed with the KniMet workflow as previously described (50). Lipids were *m/z*annotated (match to *m/z*) using the LMSD.

### Determination of 1-carbon metabolites by UHPLC-SRM-MS

The aqueous component of each extract was thoroughly dried under nitrogen and reconstituted in 25 µl aqueous 100 mM dithiothreitol (Sigma Aldrich, Haverhill, Suffolk, UK) containing 50 µM: [U^13^C, U^15^N] *L*-glutamic acid and *L*-proline (Cambridge Isotope Laboratories, Tewkesbury, MA, UK); *L*-leucine-d_10_, *L*-phenyl-d_5_-alanine and *DL*-homocysteine-3, 3, 4, 4-d_4_ (CDN Isotopes, Pointe-Claire, QC, Canada); *L*-valine-d_8_ (Sigma Aldrich) as internal standards. This mixture was thoroughly vortexed and sonicated for 15 min. at room temperature after which a further 100 µl 10 mM ammonium acetate (Sigma Aldrich) was added to the resulting solution. One-carbon cycle metabolites were analysed via a targeted assay using a Thermo Quantiva triple quadrupole mass spectrometer coupled to an Ultimate 3000 UHPLC liquid chromatography system (Thermo Scientific, Hemel Hempstead, Herts., UK). A C18PFP column (150×2.1 mm, 3 µm; Advanced Chromatography Technologies, Aberdeen, UK) was used to separate the compounds. Mobile phase A consisted of aqueous 0.1% formic acid, and mobile phase B consisted 0.1 % formic acid in acetonitrile. The following gradient was used to separate the compounds: 100% A was held for 1.5 min, followed by a linear increase of B from 0% to 100% over 3 min with a further re-equilibration for 1.5 min to give a total run time of 6 min. The flow rate was 400 µl/min and the injection volume was 2.5 µl. The mass spectrometer was operated in positive ion mode with the following parameters: a spray voltage of 3.5 kV, sheath gas 50 arb., auxiliary gas 15 arb., sweep gas 2 arbitrary units, ion transfer tube temperature 350°C and a vaporiser temperature of 400°C. All data were processed using Xcalibur (Thermo Scientific).

### Measurement of DNA Methylation

DNA was extracted using the QIAamp DNA Mini Kit (Cat. no. 51304) according to the manufacturer’s instructions but with some modifications: ∼50 mg frozen adipose tissue was pulverised in 180 µl Buffer ATL using a TissueLyser (Qiagen, West Sussex, UK) with 5 mm stainless steel beads for 20 s at 30 Hz. The sample was centrifuged briefly to ensure that all the tissue debris was on the bottom of the tube. 20 μl proteinase K was then added to the tube which was incubated for 56°C for 1 h in a shaker incubator. 200 μl Buffer AL was added to the sample, then mixed again by pulse-vortexing for 15 s, and incubated at 70°C for 10 min.

The concentration of DNA was measured by absorbance at 260 nm using a Thermo Scientific NanoDrop 2000c UV-Vis spectrophotometer. *OneStep* qMethyl™ Kit (Catalog No. D5310) was used to determine the methylation status of four genes (Lhfpl2, Rgs3, ACSL1, and AKT2). *OneStep* qMethyl™ Kit were performed on a StepOnePlus™ (Cat. No. 4376600, Thermo Scientific) using a Real-Time PCR System.

Primers used for the four genes in the Real-Time PCR processes:

ACSL1_forward, TGCGGCCGCGACTCCTTAAATA

ACSL1_reverse, AGGGAAACGAGGCCGTGGAG

AKT2_forward, CGTTGCTGCCGCCAGTTCATAAAT

AKT2_reverse, GAGCCTCCAGGTCCGTGGTC

Lhfpl2_forward, ACCGGACTGAGCGACCCTC

Lhfpl2_reverse, GGCAGGTGACAATGACATGACACATATT

Rgs3_forward, AGCCAAGTCAGGTGGAAATCT

Rgs3_reverse, CTCCATGGCGTCCCTGTT

### Measuring the ratio of methyl-cytosine to cytosine in DNA

This method is based on Rocha *et al (51)*. In brief, a calibration curve was constructed using a range of theoretical methylation percentages from 0.5 - 6.5% prepared by mixing DNA from *Escherichia coli* pBR322 DNA (New England Biolabs, N3033S; 0.5% methylation (52) and calf thymus DNA (Sigma, D4522, 6.5 % methylation (53).

25 mg of adipose tissue from mice were lysed in QIAzol reagent (Qiagen) using a tissue lyser to disrupt the tissue, and DNA extracted using the manufacturer’s guidelines. Extracted DNA was quantified using a nanodrop, by measuring absorbances at 260 nm and 280 nm. A 10 µL aliquot containing approximately 1µg of genomic DNA in a micro-vial (Agilent, cat no 22437) with 6.6 µL Internal Standard solution containing 5 nmoles [^13^C_2,_ ^15^N_2_] cytosine (Goss Scientific) and 0.16 nmoles of 5-methylcytosine (methyl-d3, 6-d1) (Qmx Laboratories Ltd) was dried in an evacuated desiccator. To hydrolyse DNA to free bases, the residue was dissolved in 20 µL of formic acid (98%), each micro-vial sealed and heated to 150 □C for 3 hours. Samples were dried overnight in an evacuated desiccator. The resulting residue was dissolved in 100 µL ammonium acetate solution (10 mM).

All analyses were performed using a Thermo scientific UHPLC+ series coupled with a TSQ Quantiva mass spectrometer (Thermo fisher scientific, Waltham, Massachusetts, United States) with an ESI source, operated in positive ion mode. The electrospray voltage was set to 3500 V for the positive ionisation and to 2500 V for the negative ionisation. Nitrogen at 48 mTorr and 420 °C was used as a drying gas for the evaporation of the eluent solution. Liquid Chromatography was performed using an ACE Excel 2 C18 PFP (100A. 150 x 2.1 mm 5µ) column. The column was conditioned at 30°C. The mobile phase consisted of: (A) 0.1% of formic acid water solution and (B) 0.1% of formic acid acetonitrile solution. The mobile phase was pumped at a flow rate of 500 µL/min programmed as follows: initially kept at 100% A for 1.60 min, then subjected to a linear decrease from 100% to 70% of A over 2.4 min and to 10% A during the next 0.5 min then kept constant for 0.5 min and brought back to initial condition after 0.1 min. The injection volume used was 8 µL. A tandem mass spectrometry fragmentation was performed to identify the analytes 5-methylcytosine (retention time = 1.1 min, positive mode, 126 → 109) and cytosine (retention time = 0.8 min, positive mode, 112 → 95). Results were expressed as 5-methylcytosine/ total-cytosine (mC/tC) ratio.

### Microarray analysis of mouse adipose tissue

RNA was extracted using the RNeasy lipid tissue kit (Qiagen GmbH, Hilden, Germany). Approximately 100 mg of tissue was used per sample for RNA isolation and procedures were carried out according to the manufacturer’s instructions. Extracted RNA was quantified by ribogreen and its purity assessed by evaluating absorbance ratios using a Fluostar microplate reader (BMG Labtech). The level of degradation of each sample was assessed using a bioanalyser (Agilent). Illumina Bead Station 500 (Illumina Inc., San Diego, CA, USA) was used to perform transcriptomics. Approximately 25,600 transcripts were interrogated using the MouseRef-8 v2.0 Expression Bead Chip, which was chosen for mouse whole-genome expression profiling.

### Data analysis methods

Multivariate statistical analysis was performed within SIMCA 14 (Umetrics, Umea, Sweden). The supervised pattern recognition tools PLS-DA, PLS, OPLS-DA and OPLS were used. PLS is a regression extension of the unsupervised pattern recognition tool principal component analysis (PCA) and finds the maximum covariance between predefined classes. The dataset is commonly visualized using scatter plots, which summarise the observations by revealing clusters and outliers in the data along the new latent dimensions in the data. Loading plots for these models display the variables (metabolites in this study) responsible for the clustering of the observations. The contribution of each variable to the clustering was ranked using the coefficient plot. The robustness of the multivariate model generated was assessed by R^2^ and Q^2^. R^2^ shows the percentage of variation explained in the model, whereas Q^2^ indicates the predictive power of the model, with Q^2^ > 30% commonly associated with robust models.

PLS-DA was used to compare different factors like genotype, age and diet with the total FAs, TGs, PCs, 1-C metabolites, transcriptomics and DNA methylation data sets. A separate model was built treating genotype, age and diet as a factor (response variable) and other omics data set as a factor (predictor variables). For each of the data types (for example: total FAs, TGs) we selected variables based on VIP scores that estimate the importance of each variable in the projection used in a PLS model. A variable with a VIP Score close to or greater than 1 can be considered important in given model (54, 55). The selection of variables was done for genotype, diet and age. O2PLS was also used to correlate between different datasets. O2PLS is a generalization of PLS and in contrast to PLS and OPLS, it is bidirectional (i.e. X ↔ Y) (56).

## Acknowledgements

This work was supported by grants from the Medical Research Council UK (MC_UP_A090_1006, MC_PC_13030, MR/P011705/1 and MR/P01836X/1) and Agilent Corporation (to JLG).

## Author Contribution

JLG and AJM conceived the original study. XW and YC performed the animal study. KL, CH, JD, LDR, MKG, and JAW performed metabolomic assays. KL, JLG, AA and RCG performed bioinformatics and interpreted the data. All authors read and approved the final manuscript.

## References

1. Adya R. Adipokines and Adipose Tissue Angiogenesis in Obesity. Immunoendocrinology. 2015.

2. Lau W.B., Ohashi K., Wang Y., Ogawa H., Murohara T., Ma X.L., et al. Role of Adipokines in Cardiovascular Disease. Circ J 2017;81(7):920–8.

3. Luo L. M. L. Adipose tissue in control of metabolism. J Endocrinol. 2016;231(3):R77–R99.

4. Booth A., Magnuson A., Fouts J. M.T. F. Adipose tissue: an endocrine organ playing a role in metabolic regulation. Horm Mol Biol Clin Investig 2016;26(1):25–42.

5. Makki K, Froguel P, Wolowczuk I. Adipose tissue in obesity-related inflammation and insulin resistance: cells, cytokines, and chemokines. ISRN Inflamm. 2013;2013:139239.

6. Sun S, Ji Y, Kersten S, Qi L. Mechanisms of inflammatory responses in obese adipose tissue. Annu Rev Nutr. 2012;32:261–86.

7. Berg AH, Scherer PE. Adipose tissue, inflammation, and cardiovascular disease. Circ Res. 2005;96(9):939–49.

8. Roberts LD, Koulman A, Griffin JL. Towards metabolic biomarkers of insulin resistance and type 2 diabetes: progress from the metabolome. Lancet Diabetes Endocrinol. 2014;2(1):65–75.

9. Hagen RM, Rodriguez-Cuenca S, Vidal-Puig A. An allostatic control of membrane lipid composition by SREBP1. FEBS Lett. 2010;584(12):2689–98.

10. Valenzuela R, Videla LA. The importance of the long-chain polyunsaturated fatty acid n-6/n-3 ratio in development of non-alcoholic fatty liver associated with obesity. Food & function. 2011;2(11):644–8.

11. Gong J, Campos H, McGarvey S, Wu Z, Goldberg R, Baylin A. Adipose tissue palmitoleic acid and obesity in humans: does it behave as a lipokine? The American journal of clinical nutrition. 2011;93(1):186–91.

12. Dennis EA, Norris PC. Eicosanoid storm in infection and inflammation. Nature reviews Immunology. 2015;15(8):511–23.

13. Borer KT. Counterregulation of insulin by leptin as key component of autonomic regulation of body weight. World journal of diabetes. 2014;5(5):606–29.

14. Shimano H. [SREBP-1c and Elovl6 as Targets for Obesity-related Disorders]. Yakugaku Zasshi. 2015;135(9):1003–9.

15. Carobbio S, Hagen RM, Lelliott CJ, Slawik M, Medina-Gomez G, Tan CY, et al. Adaptive changes of the Insig1/SREBP1/SCD1 set point help adipose tissue to cope with increased storage demands of obesity. Diabetes. 2013;62(11):3697–708.

16. Musri MM, Parrizas M. Epigenetic regulation of adipogenesis. Current opinion in clinical nutrition and metabolic care. 2012;15(4):342–9.

17. Lillycrop KA, Phillips ES, Jackson AA, Hanson MA, Burdge GC. Dietary protein restriction of pregnant rats induces and folic acid supplementation prevents epigenetic modification of hepatic gene expression in the offspring. The Journal of nutrition. 2005;135(6):1382–6.

18. Widiker S, Karst S, Wagener A, Brockmann GA. High-fat diet leads to a decreased methylation of the Mc4r gene in the obese BFMI and the lean B6 mouse lines. Journal of applied genetics. 2010;51(2):193–7.

19. Zeisel SH. Metabolic crosstalk between choline/1-carbon metabolism and energy homeostasis. Clin Chem Lab Med. 2013;51(3):467–75.

20. Corbin KD, Zeisel SH. Choline metabolism provides novel insights into nonalcoholic fatty liver disease and its progression. Curr Opin Gastroenterol. 2012;28(2):159–65.

21. Anderson OS, Sant KE, Dolinoy DC. Nutrition and epigenetics: an interplay of dietary methyl donors, one-carbon metabolism and DNA methylation. J Nutr Biochem. 2012;23(8):853–9.

22. Noureddin M, Mato JM, Lu SC. Nonalcoholic fatty liver disease: update on pathogenesis, diagnosis, treatment and the role of S-adenosylmethionine. Experimental biology and medicine. 2015;240(6):809–20.

23. Sud M, Fahy, E., Cotter, D., Brown A., Dennis, E., Glass, C., Murphy, R., Raetz, C., Russell, D., Subramaniam, S. LMSD: LIPID MAPS structure database. Nucleic Acid Research. 2006;35:D527–32.

24. Multhaup ML, Seldin MM, Jaffe AE, Lei X, Kirchner H, Mondal P, et al. Mouse-human experimental epigenetic analysis unmasks dietary targets and genetic liability for diabetic phenotypes. Cell metabolism. 2015;21(1):138–49.

25. Benton MC, Johnstone A, Eccles D, Harmon B, Hayes MT, Lea RA, et al. An analysis of DNA methylation in human adipose tissue reveals differential modification of obesity genes before and after gastric bypass and weight loss. Genome biology. 2015;16:8.

26. Dick KJ, Nelson CP, Tsaprouni L, Sandling JK, Aissi D, Wahl S, et al. DNA methylation and body-mass index: a genome-wide analysis. Lancet. 2014;383(9933):1990–8.

27. Ramanadham S, Zhang S, Ma Z, Wohltmann M, Bohrer A, Hsu FF, et al. Delta6-, Stearoyl CoA-, and Delta5-desaturase enzymes are expressed in beta-cells and are altered by increases in exogenous PUFA concentrations. Biochimica et biophysica acta. 2002;1580(1):40–56.

28. Park HG, Kothapalli KS, Park WJ, DeAllie C, Liu L, Liang A, et al. Palmitic acid (16:0) competes with omega-6 linoleic and omega-3 a-linolenic acids for FADS2 mediated Delta6-desaturation. Biochimica et biophysica acta. 2016;1861(2):91–7.

29. Lopez-Vicario C, Gonzalez-Periz A, Rius B, Moran-Salvador E, Garcia-Alonso V, Lozano JJ, et al. Molecular interplay between Delta5/Delta6 desaturases and long-chain fatty acids in the pathogenesis of non-alcoholic steatohepatitis. Gut. 2014;63(2):344–55.

30. Kien CL, Bunn JY, Poynter ME, Stevens R, Bain J, Ikayeva O, et al. A lipidomics analysis of the relationship between dietary fatty acid composition and insulin sensitivity in young adults. Diabetes. 2013;62(4):1054–63.

31. Perez-Jimenez F, Lopez-Miranda J, Pinillos MD, Gomez P, Paz-Rojas E, Montilla P, et al. A Mediterranean and a high-carbohydrate diet improve glucose metabolism in healthy young persons. Diabetologia. 2001;44(11):2038–43.

32. Sjogren P, Sierra-Johnson J, Gertow K, Rosell M, Vessby B, de Faire U, et al. Fatty acid desaturases in human adipose tissue: relationships between gene expression, desaturation indexes and insulin resistance. Diabetologia. 2008;51(2):328–35.

33. Yew Tan C, Virtue S, Murfitt S, Robert LD, Phua YH, Dale M, et al. Adipose tissue fatty acid chain length and mono-unsaturation increases with obesity and insulin resistance. Sci Rep. 2015;5:18366.

34. Pietilainen KH, Rog T, Seppanen-Laakso T, Virtue S, Gopalacharyulu P, Tang J, et al. Association of lipidome remodeling in the adipocyte membrane with acquired obesity in humans. PLoS Biol. 2011;9(6):e1000623.

35. Lopategi A, Lopez-Vicario C, Alcaraz-Quiles J, Garcia-Alonso V, Rius B, Titos E, et al. Role of bioactive lipid mediators in obese adipose tissue inflammation and endocrine dysfunction. Mol Cell Endocrinol. 2015.

36. Jump DB. Dietary polyunsaturated fatty acids and regulation of gene transcription. Current opinion in lipidology. 2002;13(2):155–64.

37. Wang X, West JA, Murray AJ, Griffin JL. Comprehensive Metabolic Profiling of Age-Related Mitochondrial Dysfunction in the High-Fat-Fed ob/ob Mouse Heart. J Proteome Res. 2015;14(7):2849–62.

38. Fagone P, Jackowski S. Phosphatidylcholine and the CDP-choline cycle. Biochimica et biophysica acta. 2013;1831(3):523–32.

39. Esfandiari F, You M, Villanueva JA, Wong DH, French SW, Halsted CH. S-adenosylmethionine attenuates hepatic lipid synthesis in micropigs fed ethanol with a folate-deficient diet. Alcoholism, clinical and experimental research. 2007;31(7):1231–9.

40. Christensen KE, Mikael LG, Leung KY, Levesque N, Deng L, Wu Q, et al. High folic acid consumption leads to pseudo-MTHFR deficiency, altered lipid metabolism, and liver injury in mice. The American journal of clinical nutrition. 2015;101(3):646–58.

41. Cummins TD, Holden CR, Sansbury BE, Gibb AA, Shah J, Zafar N, et al. Metabolic remodeling of white adipose tissue in obesity. Am J Physiol Endocrinol Metab. 2014;307(3):E262–77.

42. Sun K, Kusminski CM, Scherer PE. Adipose tissue remodeling and obesity. J Clin Invest. 2011;121(6):2094–101.

43. Lee MJ, Wu Y, Fried SK. Adipose tissue remodeling in pathophysiology of obesity. Current opinion in clinical nutrition and metabolic care. 2010;13(4):371–6.

44. Shimano H. SREBPs: physiology and pathophysiology of the SREBP family. The FEBS journal. 2009;276(3):616–21.

45. Roberts LD, Murray AJ, Menassa D, Ashmore T, Nicholls AW, Griffin JL. The contrasting roles of PPARdelta and PPARgamma in regulating the metabolic switch between oxidation and storage of fats in white adipose tissue. Genome biology. 2011;12(8):R75.

46. Acharjee A, Ament, Z., West, J.A., Stanley, E., Griffin, J.L. Integration of metabolomics, lipidomics and clinical data using a machine learning method. BMC Bioinformatics. 2016;17:440.

47. Murray AJ, Knight NS, Cochlin LE, McAleese S, Deacon RM, Rawlins JNP, et al. Deterioration of physical performance and cognitive function in rats with short-term high-fat feeding. The FASEB Journal. 2009;23(12):4353–60.

48. Folch J, Lees M, Sloane-Stanley G. A simple method for the isolation and purification of total lipids from animal tissues. J biol chem. 1957;226(1):497–509.

49. Bligh EG, Dyer WJ. A rapid method of total lipid extraction and purification. Canadian journal of biochemistry and physiology. 1959;37(8):911–7.

50. Liggi S., Hinz C., Hall Z., Santoru M.L., Poddighe S., Fjeldsted J., et al. KniMet: a pipeline for the processing of chromatography-mass spectrometry metabolomics data. Metabolomics. 2018;14(4):52.

51. Rocha M, Castro R, Rivera I. Global DNA methylation: comparison of enzymatic- and non-enzymatic-based methods.. Clinical Chemistry and Laboratory Medicine. 2010;48(12):1793–179.

52. Eick D, Fritz H, Doerfler W. Quantitative determination of 5-methylcytosine in DNA by reverse-phase high performance liquid chromatography. Anal Biochem. 1983;135:165–71.

53. Kok R, Smith D, Barto R. Global DNA methylation measured by liquid chromatography-tandem mass spectrometry: analytical technique, reference values and determinants in healthy subjects. Clinical Chemistry and Laboratory Medicine. 2007;45(7):903–11.

54. Galindo-Prieto B, Eriksson, L., Trygg, J. Variable influence on projection (VIP) for OPLS models and its applicability in multivariate time series analysis. Chemometr Intell Lab. 2015;146:297–304.

55. Palermo G, Piraino, P., Zucht, H.D. Performance of PLS regression coefficients in selecting variables for each response of a multivariate PLS for omics-type data. Adv Appl Bioinform Chem. 2009;2:57–70.

56. Bylesjo M, Eriksson D, Kusano M, Moritz T, Trygg J. Data integration in plant biology: the O2PLS method for combined modeling of transcript and metabolite data. Plant J. 2007;52(6):1181–91.

